# Influence of temperature on the development, reproduction and regeneration in the flatworm model organism *Macrostomum lignano*

**DOI:** 10.1101/389478

**Authors:** Jakub Wudarski, Kirill Ustyantsev, Lisa Glazenburg, Eugene Berezikov

**Affiliations:** European Research Institute for the Biology of Ageing, University of Groningen, University Medical Center Groningen, Antonius Deusinglaan 1, 9713AV, Groningen, The Netherlands; Institute of Cytology and Genetics, Prospekt Lavrentyeva 10, 630090, Novosibirsk, Russia.

**Keywords:** Temperature, Heat Shock Response, Flatworms, *Macrostomum lignano*, Fertility, Regeneration.

## Abstract

The free-living marine flatworm *Macrostomum lignano* is a powerful model organism to study mechanisms of regeneration and stem cell regulation due to its convenient combination of biological and experimental properties, including the availability of transgenesis methods, which is unique among flatworm models. However, due to its relatively recent introduction in research, there are still many biological aspects of the animal that are not known. One of such questions is the influence of the culturing temperature on *Macrostomum* biology. Here we systematically investigated how different culturing temperatures affect the development time, reproduction rate, regeneration, heat shock response, and gene knockdown efficiency by RNA interference in *M. lignano.* We used marker transgenic lines of the flatworm to accurately measure the regeneration endpoint and to establish the stress response threshold for temperature shock. We found that compared to the culturing temperature of 20°C commonly used for *M. lignano*, elevated temperatures of 25°C-30°C substantially speed-up the development and regeneration time and increase reproduction rate without detectable negative consequences for the animal, while temperatures above 30°C elicit a heat shock response.

We show that altering the temperature conditions can be used to shorten the time required to establish *M. lignano* cultures, store important lines and optimize the microinjection procedures for transgenesis. Our findings will help to optimize the design of experiments in *M. lignano* and thus facilitate future research in this model organism.

## Background

Flatworms (Platyhelminthes) are a large phylum in the animal kingdom (Metazoa) with a broadly present capacity to regenerate lost tissues and body parts among its representatives [1]. This regenerative ability attracted the interests of scientists for a long time, and in particular the free-living planarian flatworms (Tricladida) *Schmidtea mediterranea* and *Dugesia japonica* have been studied extensively and yielded numerous insights into the mechanisms of regeneration [2–5]. More recently, a non-planarian flatworm model *Macrostomum lignano* (Macrostopmorpha) has entered the regeneration research arena, bringing an attractive combination of experimental and biological features [6,7]. *M. lignano* is a free-living marine flatworm capable of regeneration anterior to the brain and posterior to the pharynx [8]. Similar to other flatworms, regeneration in *M. lignano* is fueled by stem cells called neoblasts [9]. It is a small and transparent animal that is easy to culture in laboratory conditions. These features, together with the recently reported genome and transcriptome assemblies [10,11] and the development of a robust transgenesis method [12] make *M. lignano* a versatile model organism for research on stem cells and regeneration [7].

*M. lignano* is a non-self-fertilizing hermaphrodite with a short generation time of 2-3 weeks [13]. When cultured in standard laboratory conditions, animals lay approximately one egg per day. Embryonic development takes 5 days, and hatchlings reach adulthood in about two weeks. The laid eggs are fertilized, relatively large (100 µm) and follow the archoophoran mode of development [13], i.e. they have a large, yolk-rich oocyte instead of separate yolk cells that supply a small oocyte. These properties of the eggs make them a good target for delivery of external agents, such as DNA, RNA and protein, by the means of microinjections.

The possibility to introduce foreign genetic material and modify the genome of an animal is a highly sought-after property and an integral part of the genomic toolkit in model organisms commonly used for genetic research, such as the nematode *Caenorhabditis elegans*, fruitfly *Drosophila melanogaster*, yeast and mouse, since it broadens the experimental approaches and greatly improves the chances to decipher biological phenomena that scientists want to study. We have recently demonstrated that microinjection of DNA into single-cell stage embryos can be used to generate transgenic *M. lignano* animals [12]. The technique is efficient and robust, and stable transgenic lines can be obtained within three-to-four months, including F1 and F2 crosses. However, from the experimental perspective it would be advantageous if the time required to generate transgenic animals can be shortened further, and manipulation of culture temperature conditions is a way to approach this.

Temperature is one of the most important factors for most biological processes. The overall range of temperatures that support active life on Earth stretches from as low as -1.8°C in polar regions to around 113°C for thermophilic archaea that stand on the other side of the extreme [14,15]. Most animals have their specific temperature range in which their growth is optimal to them; this is because even small alterations in the temperature can lead to great changes in metabolism of the animals. If these changes are sustainable for the organism, the increase in the temperature usually leads to an increase in the speed of physiological processes. The most common way of showing this relationship is using the temperature coefficient Q10 = (Rate 2 / Rate 1)^10/(Temperature^ ^2−Temperature^ ^1)^, which compares the rates of a process at a given temperature with the rate at temperature increased by 10°C [16]. The most popular way of using the Q10 is by measuring the oxygen consumption [17], but it may also be applied to various different measurements such as electric organ discharge [18] or locomotor performance [19].

All living organisms respond to changes in the temperature of their environment. Perturbations in the temperature will usually trigger the heat shock response pathway. It is an ancient and universal mechanism based on specialized chaperone molecules called heat shock proteins, or Hsps [20]. These proteins can help other proteins to fold correctly, repair damaged proteins or degrade them. They can also be a good indicator of stress that an organism undergoes [21].

Here we present how temperature affects development, growth, fertility and regeneration capabilities of *Macrostomum lignano*. We have tested the stress response to elevated temperatures by measuring the activity of the Heat shock 20 (*Hsp20*) gene using qRT-PCR and *Hsp20::mScarlet* transgene expression. Furthermore, we have measured the hatching speed of the eggs when incubated at different temperatures, as well as the number of offspring produced in these conditions. We also investigated how changes in temperature influence the regeneration speed of the animals, and the efficiency of gene knockdown by RNA interference. Out findings establish optimal conditions for *M. lignano* cultures and will instruct future research in this model organism, particularly for creating transgenic animals.

## Methods

### Strains

The DV1 inbred *M. lignano* line used in this study was described previously [10,22,23]. The NL10 and NL22 lines were previously established in our laboratory [12]. Animals were cultured under laboratory conditions in plastic Petri dishes (Greiner), filled with nutrient enriched artificial sea water (Guillard’s f/2 medium). Worms were fed ad libitum on the unicellular diatom *Nitzschia curvilineata* (Heterokontophyta, Bacillariophyceae) (SAG, Göttingen, Germany). The conditions in the climate chambers were set at 20°C, 25°C, 30°C and 35°C with constant aeration, and a 14h/10h day/night cycle.

The heat shock sensor construct KU#49 was created using a previously described double-promoter vector approach [12]. The promoter region of *M. lignano hsp20* homolog gene (Mlig005128.g2) was cloned using primers 5’-GGATGGATCCTCATTTATAAGCGTACCGTACT-3’ and 5’-TTATAAGCTTCATGCTGTTGTTGACTGGCGTA-3’ to drive expression of mScarlet-I red fluorescent protein, while elongation factor 1 alpha (*EFA*) promoter driving expression of NeonGreen fluorescent protein was used for the selection of transgenic animals. 235 single-cell eggs were injected with the KU#49 plasmid as previously described [12] but without radiation treatment, the hatchlings were selected based on the presence of green fluorescence and a stable transgenic line NL28 was established

### Egg hatching

Twenty single-cell stage eggs per temperature condition were picked and transferred to single wells in a 6 well plate. They were monitored daily and hatched worms were immediately removed from the test well. Each temperature condition was tested independently in triplicate.

### Heat shock

For measuring expression level of the *Mlig-hsp20* gene by qRT-PCR, 50 worms of the same age were selected for each of the three replicates. Animals were incubated for 2 hours at different temperatures (20°C, 25°C, 33°C, 34°C and 35°C) and 2 more hours at 20°C, before the RNA extraction (RNeasy, Qiagen). The quantitative PCR was done using the Light Cycler 480 (Roche) with 5’-CGAAGATGTCACTGAGGTCAAG-3’ and 5’-GCGCCTGCAGTAGAAGAAT-3’ as primers and GAPDH, COX and EIF as reference target genes as previously described [24]. The analysis of the results was performed using the qBase+ software (Biogazelle).

For monitoring heat shock response using *Hsp20::mScrarlet* transgene, NL28 transgenic animals were incubated for 2 hours at different temperatures (20°C, 25°C to 35°C with 1°C interval, and 37°C), followed by 22 hours at 20°C, and then imaged using Zeiss Axio Zoom V16 microscope with an HRm digital camera and Zeiss filter sets 38HE (FITC) and 43HE (DsRed). The pictures were analyzed using ImageJ software. Images were converted to 8-bits, the area of the worm was selected and the median value of the signal was measured and visualized using box plot. Graphs were made using the ggplot2 package for R.

### Reproduction

Three sets of 20 freshly hatched worms were selected per line and temperature condition. They were kept in the selected temperature until the end of the experiment, which lasted for 5 weeks. Twice per week the worms were transferred to new plates with fresh food, and the old plates, where animals laid eggs in the preceding time interval, were incubated until all eggs were hatched, after which the hatchlings were counted. The approach of counting hatchlings as a proxy to the number of laid eggs rather than the direct counting of eggs was chosen because it is easier and more reliable to count hatchlings compared to the egg clumps.

### Regeneration

Twelve NL22 worms expressing GFP marker in testes [12] were used per condition and cut above the testes. The worms were then placed in 12-well plates with fresh diatom and monitored daily. The days of the first appearance of GFP signal in the testes and in the seminal vesicle were noted and used to measure the time required for regeneration.

### RNA interference

Fifteen worms were selected per temperature condition (20°C, 25°C and 30°C) and were treated with *Mlig-ddx39* dsRNA fragments as previously described [11]. The morphology and viability of the worms was monitored on a daily basis and any abnormalities were noted. GFP dsRNA was used as a negative control.

## Results

### Establishing the temperature range

Commonly used laboratory conditions for *Macrostomum lignano* cultures are as follows: temperature of 20°C, humidity 60% and light/dark cycle of 14h/10h. These conditions were chosen mainly because they are optimal for the growth of the diatom *Nitzschia curvilineata*, which is the main food source for the worms [25]. To assess the temperature conditions that can be used in the experiment, we have first established the temperature range in which the worms survive. While freezing the worms proved to be lethal, they could survive when kept at 4°C for at least two weeks. However, because the diatom does not grow in these conditions, we decided to exclude 4°C from further experiments. Other temperatures below 20°C were also excluded from the experiment, since the primary objective of the study was to find conditions that accelerate growth and development. On the other side of the temperature spectrum the worms dissolve when kept at 42°C for 2 hours, and died after one week of culture at 37°C. Therefore, we have decided to use 20°C, 25°C, 30°C and 35°C as our experimental conditions to study long-term temperature effects on *M. lignano* (Fig. 1).

**Figure 1.**
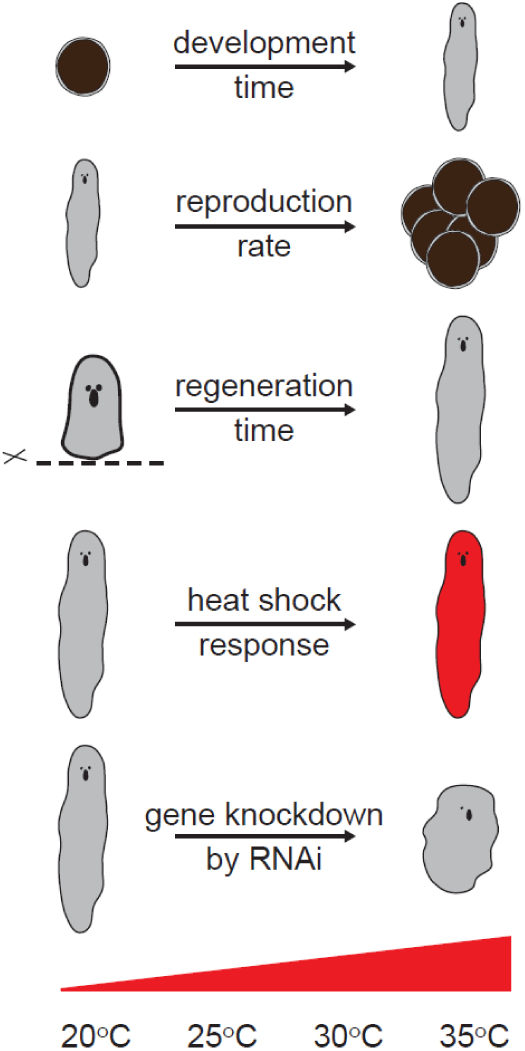
Design of the study Embryos and animals of two wild-type strains, DV1 and NL10, were cultured at a range of temperatures from 20°C to 35°C and development time, reproduction, regeneration time, heat shock response and efficiency of gene knockdown by RNA interference were measured

### Heat shock response

To investigate which temperatures induce stress response in the worms, we monitored the activity of the heat shock 20 promoter (*Mlig*-*hsp20*). First, we performed a quantitative RT-PCR to measure the expression level of *Mlig-hsp20* at 20°C, 25°C, 30°C, 33°C, 34°C and 35°C. There was no significant difference in the expression level of Hsp20 between 20°C and 25°C (Fig. 2a). However, a small (2-fold) but significant (P = 0.027, *t-*test) increase in the expression was observed at 30°C compared to 20°C (Fig. 2a). More than a ten-fold raise in the expression level was observed at 33°C, which increased to more than 100-fold at the highest tested temperature of 35°C (Fig. 2a). In the second test we have created a transgenic line expressing mScarlet-I protein under the control of the *Mlig-hsp20* promoter and measured the level of fluorescence 24 hours after a 2 hour incubation at different temperatures ranging from 20°C to 37°C (Fig. 2b,c). More than two- and ten-fold increase in the fluorescence was observed at 34°C and 37°C respectively (Fig. 2b).

**Figure 2.**
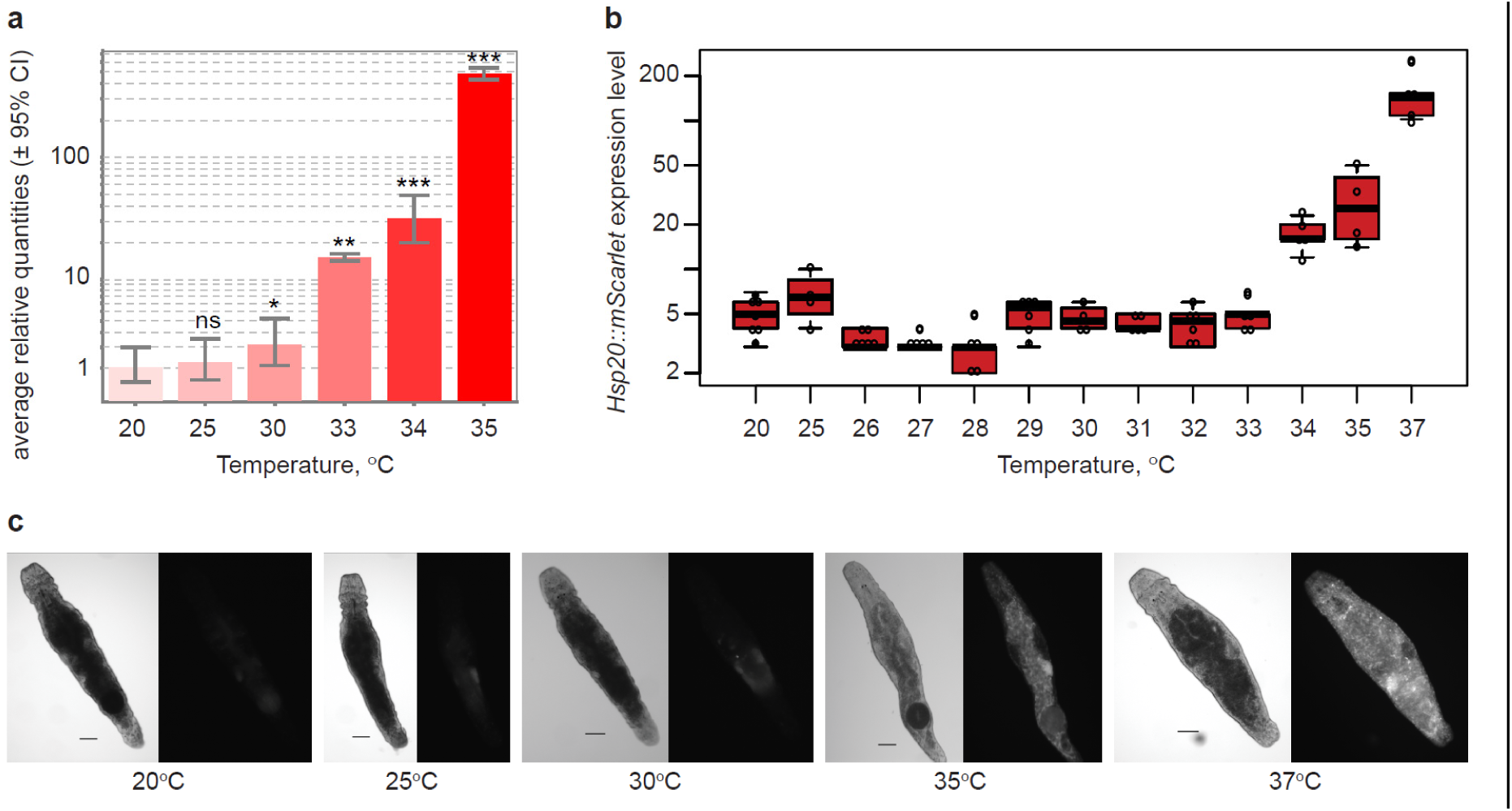
Heat shock response in *M. lignano*. (a) RT-qPCR analysis of *Mlig-hsp20* gene expression at different temperatures. The graph is normalized to the expression levels at 20°C. Statistical significance of the changes relative to the 20°C condition is calculated using *t*-test. ns, P > 0.05; *, P ≤ 0.05; **, P ≤ 0.01; ***, P ≤ 0.001. Error bars indicate standard deviation. (b) Fluorescence intensity of *hsp20::mScarlet* transgene expression at different temperatures. Error bars indicate standard deviation. (c) Example images of NL28 transgenic animals used for measuring *hsp20::mScarlet* transgene expression. DIC and dsRed channels are shown for each temperature. Scale bars are 100 µm.

### Speed of embryonic development

According to Morris et al. [13], it takes around 120h (5 days) for *Macrostomum* eggs to fully develop when eggs are kept at 20°C. To investigate how the temperature affects the speed of embryonic development and hatching, we picked freshly laid embryos and monitored their development until hatching at different temperatures. To investigate potential differences due to genetic background, we have used two *M. lignano* lines independently derived from wild-type populations, the DV1 and NL10 lines, which differ by a whole-chromosome duplication [12]. We first studied the effect of low temperature. At 4°C the development of the eggs is arrested and they can be stored for at least one month, and resume their development once returned to higher temperatures. Next we studied how quickly the eggs will develop at temperatures between 20°C and 35°C. As shown in Fig. 3, when kept at standard conditions (20°C) the eggs started to hatch after 6 days. Increase in the temperature resulted in proportionally faster embryonic development and earlier hatching, which was two times faster at 35°C compared to 20°C and took only 3 days. Of note, about 10% of eggs are still non hatched after 8 days of incubation at 20°C, compared to less than 5% at higher temperatures, suggesting that even the highest tested temperature of 35°C does not have detrimental effects on the survival of embryos.

**Figure 3.**
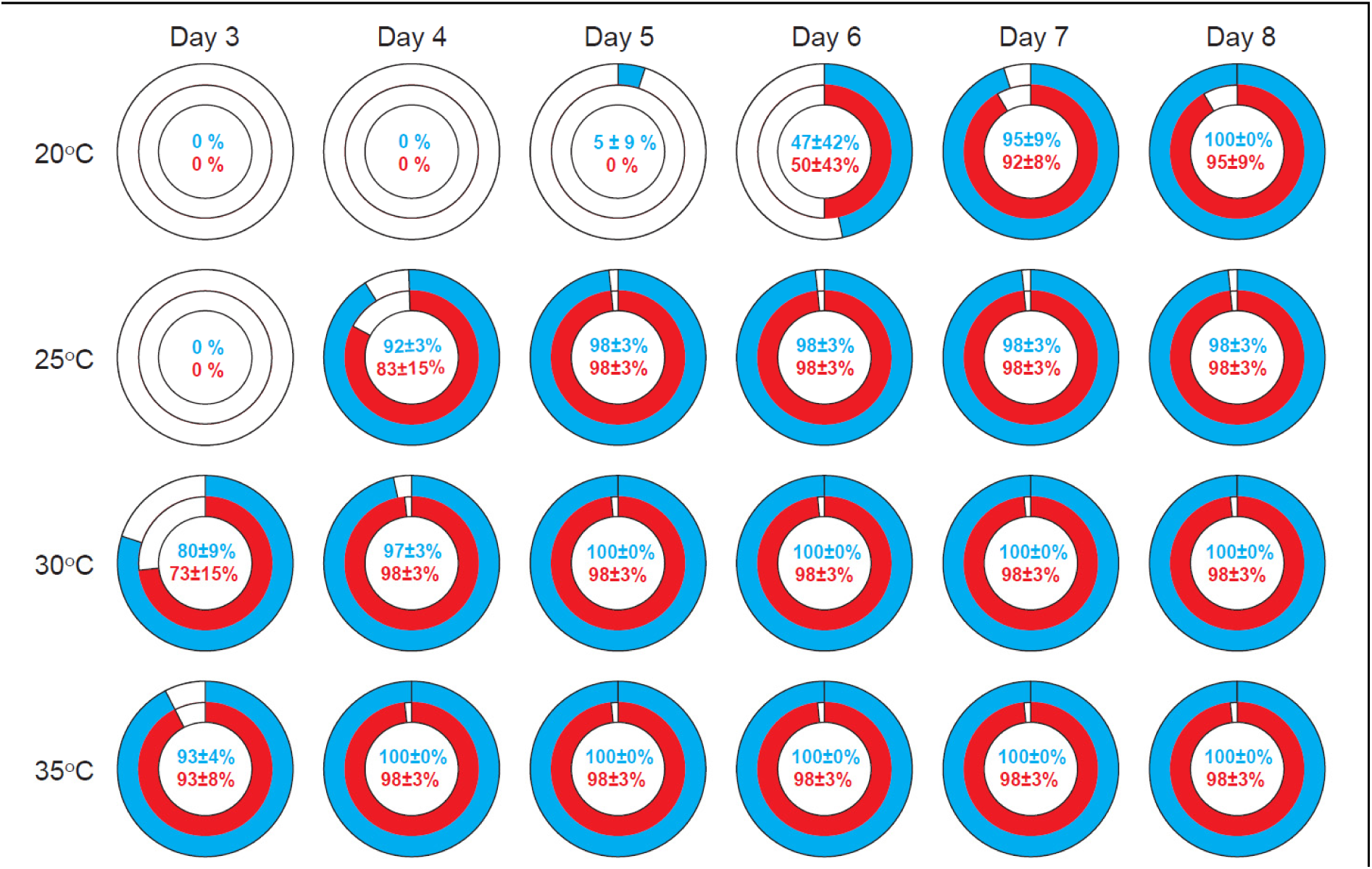
Time of embryonic development of *M. lignano* at different incubation temperatures. DV1 (red) and NL10 (blue) *M. lignano* lines were used in the experiment and 20 eggs were monitored per conditions. The experiment was repeated three times. Mean ± standard deviation values are indicated.

### Reproduction

The reproduction rate is a very important factor for a model organism, since animals with shorter generation time enable faster generation of data in genetic experiments. In addition, if animals produce large numbers of offspring, the generated data will, in most cases, have higher statistical power.

To assess the temperature impact on *M. lignano* reproduction rate we have compared the number of offspring generated along the course of five weeks by worms kept at different temperature conditions. The experiment was started with hatchlings to incorporate postembryonic development in the study, and both DV1 and NL10 line were used. It took the hatchlings of both lines three weeks to grow and produce first progeny at 20°C, while at 25°C hatchlings were observed in two weeks for NL10 line but not for DV1. At 30°C and 35°C both lines produced progeny already after two weeks (Fig. 4). From three weeks onwards the number of hatchlings produced per week increased from below 200 at 20°C to more than 300 in temperatures above 20°C and was the highest for the worms kept at 30°C. This was true for both genetic backgrounds, and we did not observe significant differences between DV1 and NL10 lines (Fig. 4).

**Figure 4.**
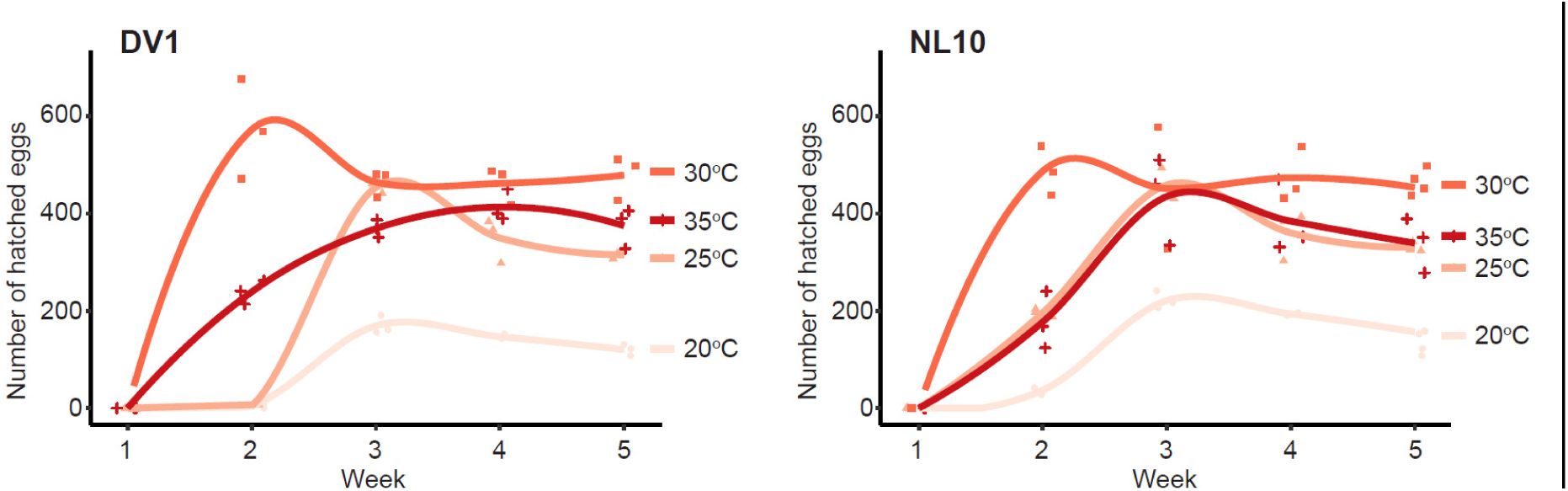
Effect of incubation temperature on reproduction rate in DV1 and NL10 *M. lignano* lines. For each condition 20 hatchlings were selected and their hatched eggs counted every week. Experiments were performed in triplicate.

Although temperatures of 30°C and higher resulted in more hatchlings, it also activates a heat shock response (Fig. 2). To investigate the long-term effect of the elevated temperature and potential stresses that it can have on the worms, we kept the worms at the selected temperatures and monitored for morphological aberrations after three and six months of culturing. Worms kept at 25°C showed no morphological changes at both check points as compared to the worms kept at 20°C (Fig. 5). However, for both the 30°C and 35°C temperature conditions, various aberrations in the general morphology were observed. Already after three months an increased number of cysts, damaged tissue and enlarged testes were commonly present in both DV1 and NL10 lines (Fig. 5). None of the worms kept at 35°C survived till the six months checkpoint, and all worms that survived at 30°C showed morphological aberrations (Fig. 5).Therefore, prolonged exposure to the temperatures above 30°C is detrimental for *M. lignano*.

**Figure 5.**
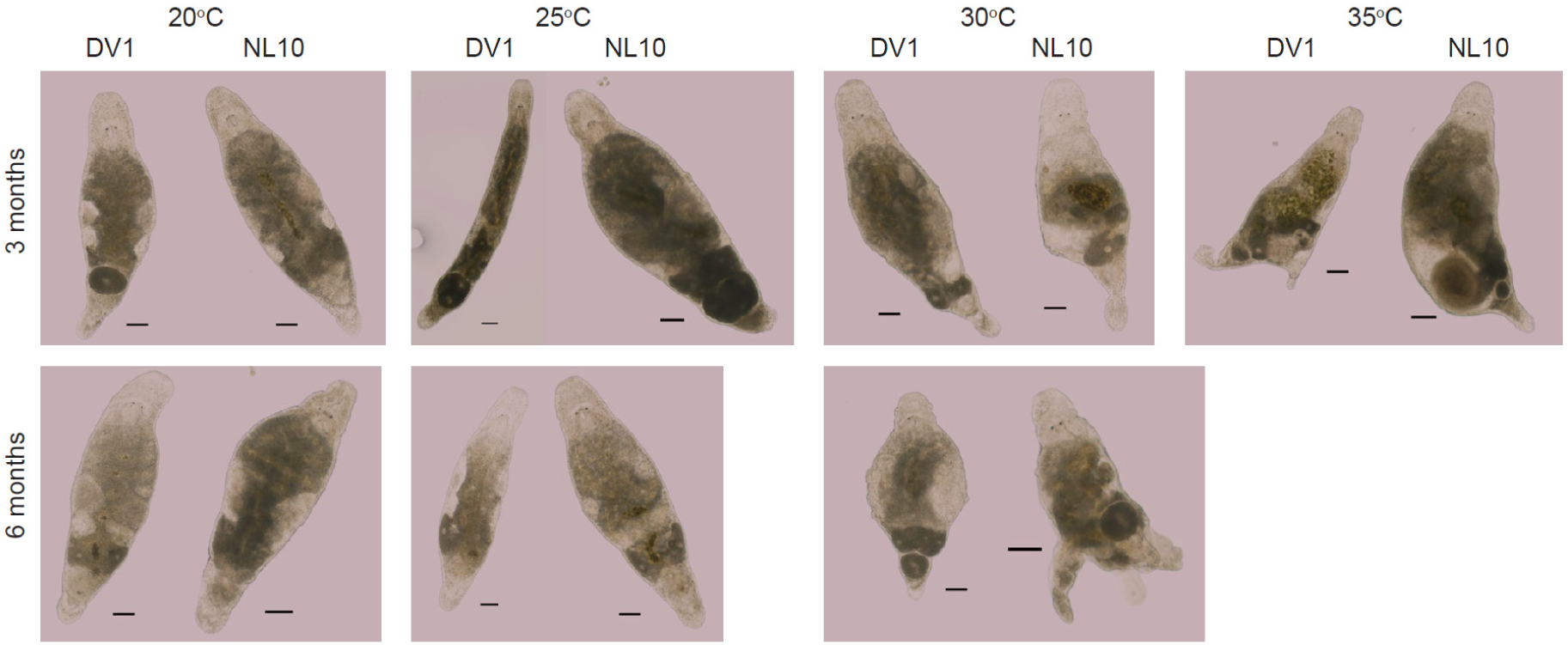
Effects of prolonged exposure to high temperatures on morphology in *M. lignano*. No visible abnormalities after incubation at 20°C or 25°C for up to 6 months.

### Regeneration time

Since the main attraction of *M. lignano* as a model organism is its regenerative ability, we next studied how temperature influences regeneration. It is commonly accepted that the increase of temperature leads to the overall metabolism increases [16]. To evaluate the influence of the temperature on the time needed for a worm to fully regenerate its body after amputation above the testes region, we tracked the regeneration of testes and the appearance of sperm in the vesicular seminalis using a previously established transgenic line NL22, which expresses GFP under the control of the testes- and sperm-specific ELAV promoter [12]. Using such a transgenic marker provides better precision, consistency and efficiency in assaying the extent of regeneration. Indeed, we observed no significant variation when assessing the time of appearance of GFP signal after amputation at all tested temperatures (Table 1).

**Table 1.** Effect of temperature on the speed of regeneration

| Temperature | Regeneration time, days* |  |
| --- | --- | --- |
|  | Testes | Sperm |
| 20°C | 6 | 8 |
| 25°C | 3 | 5 |
| 30°C | 2 | 3 |
| 35°C | 2 | 3 |

\* no variation was observed, standard deviation = 0 in all conditions.

As expected, the speed of regeneration increased with temperature (Table 1). If we take the regeneration time at 20°C as the standard rate, then for testes regeneration the calculated temperature coefficients are Q10=4 at 25°C and Q10=3 for 30°C and 35°C. This shows that the largest effect is obtained when increasing the temperature to 25°C, and the fastest regeneration takes place at 30°C, without further benefit of increased temperature on the regeneration time. Hence, temperatures of 25°C and 30°C ought to be considered in regeneration experiments on *M. lignano*, as it shortens the duration of an experiment by two-to-three times (Table 1).

### RNA interference

Knockdown of gene expression by RNA interference (RNAi) is currently the primary approach for loss-of-function studies in *M. lignano* [7]. In this approach animals are soaked with double-stranded RNA against the target gene, and often prolonged treatments for several weeks are required to observe a phenotype [11,26,27]. We have tested how temperature influences the speed of phenotype development upon RNAi treatment. To do this we knocked down *Mlig-ddx39*, a gene known for its function in cell proliferation *in M. lignano* and a robust lethal knockdown phenotype [11]. Similarly to the reproduction test, we used the DV1 and NL10 lines for this experiment (Table 2). At 20°C it took around 20 days for all animals to die upon *Mlig-ddx39* knockdown. When the worms were kept at higher temperatures, the death occurred faster: at 11 and 8 days for the worms kept at 25°C and 30°C respectively. There was no visible difference between the two strains used for the experiment (Table 2). Hence, similar to regeneration, RNAi is scaled with temperature in *M. lignano* and higher temperatures can be used to shorten the duration of experiments.

**Table 2.** Effect of temperature on gene knockdown by RNAi

| Temperature | Incubation time with dsRNA against *Mlig-ddx39* gene until lethality*, days, mean ± standard deviation | |
| --- | --- | --- |
|  | DV1 | NL10 |
| 20°C | 17.8±1.3 | 16.6±1.9 |
| 25°C | 10.1±0.6 | 9.5±0.5 |
| 30°C | 7.9±0.2 | 7.2±0.4 |

\* no lethality was observed at the tested temperatures when incubating animals with dsRNA against *gfp* as a control.

## Discussion

Temperature is one of the key aspects in husbandry of laboratory animals, and knowing the optimal values helps to provide them with the most suitable conditions. Changing the temperature has long been used as a method to influence the growth of model organisms, as best seen in the case of *C. elegans* [28,29].

Here we tested how different temperatures affect the biology of the flatworm *Macrostomum lignano*. While worms can be kept at 4°C for two weeks, the freshly laid eggs can be stored in these conditions for much longer. Practically, this is very useful for back-up and long-term storage of valuable worm cultures, such as transgenic lines, and for collection and synchronization of eggs for subsequence microinjection. The possibility to collect one-cell stage eggs and prevent their division by keeping them on ice or in a fridge provides a wider time window for microinjections, since it allows splitting the egg picking from their injections. At the same time, higher temperatures speed-up the development and shorten the time required for the worms to hatch. Incubating the eggs in 25°C or 30°C can save several days of unnecessary waiting time, enabling faster generation of transgenic animals.

Standard *M. lignano* cultures are kept in 20°C and are transferred to fresh diatom every week, or twice a week when cultures are used for egg production for microinjections [12]. In this way, a new population of worms can be expanded every two weeks if the starting culture is sufficiently dense. The generation time of two weeks can be quite limiting when experiments require large numbers of specimens or when the starting culture has low number of animals. This is often the case for establishing a new transgenic line, performing *in situ* hybridization, RNAi experiment or isolating nucleic acids and proteins. Increasing overall egg production by simply putting the cultures at higher temperature can be used as an easy method to quickly generate large number of worms. However, one must be cautious to avoid undesirable stress response that could potentially lead to distorted results. While keeping the animals at 30°C can result in a significant increase in the egg production, the changes in the morphology, visible already after 3 months, indicate that these conditions might cause too much stress to the worms. Indeed, we observed mild heat shock response activation at 30°C, which quickly increases with further rising of incubation temperature.

Regeneration is one of the most prominent features of the flatworms, and *M. lignano* provides a powerful experimental platform to study this phenomenon. We demonstrate that speed of regeneration increases with temperature in *M. lignano*, which can be taken into account and used to shorten time required for an experiment when needed. Similarly, gene knockdown by RNA interference is temperature-dependent and RNAi experiments can be accelerated by increasing the temperature.

In this work we show that simple temperature control can significantly benefit a wide range of experiments using the flatworm model organism *Macrostomum lignano*. Based on our results and experience, we propose the following: (1) use 4°C for storing the eggs and keeping them prior to microinjections; (2) use 20°C for standard cultures that do not need rapid expansion, thus reducing the frequency of transferring the animals to a fresh food source; (3) use 25°C for cultures that need to be quickly expanded and for the eggs that need to be hatched faster, for example microinjected eggs; (4) use 25°C or 30°C for RNAi experiments to observe the desired phenotype faster.

## Acknowledgements

We thank Stijn Mouton for valuable comments on the manuscript. The DV1 line used in this study is a gift from Schärer laboratory.

## Author contributions

JW, KU, LG and EB designed the study. JW, KU and LG performed experiments. JW, LG and EB analyzed the results. JW and EB wrote the manuscripts. All authors read and approved the final manuscripts.

